# Improving the sensitivity of in vivo CRISPR off-target detection with DISCOVER-Seq+

**DOI:** 10.1101/2022.10.29.514376

**Authors:** Roger S. Zou, Yang Liu, Oscar E. Reyes Gaido, Maximilian F. Konig, Brian J. Mog, Leo L. Shen, Franklin Aviles-Vazquez, Alberto Marin-Gonzalez, Taekjip Ha

## Abstract

Discovery of off-target CRISPR-Cas genome editing activity in patient-derived cells and animal models is crucial for therapeutic applications, but currently exhibits low sensitivity. We demonstrate that inhibition of DNA-dependent protein kinase catalytic subunit (DNA-PKcs) accumulates repair protein MRE11 at CRISPR-targeted sites, enabling high-sensitivity mapping of off-target sites to positions of MRE11 binding using chromatin immunoprecipitation sequencing (ChIP-seq). This technique, termed DISCOVER-Seq+, discovered up to 5-fold more CRISPR off-target sites in immortalized cell lines, primary human cells, and mice compared to previous methods. We demonstrated applicability to ex vivo knock-in of a cancer-directed transgenic T-cell receptor in primary human T cells and in vivo adenovirus knock-out of cardiovascular risk gene *PCSK9* in mice. DISCOVER-Seq+ is the most sensitive method to-date for discovering off-target genome editing in vivo.

## MAIN TEXT

CRISPR-Cas genome editing is a transformative technology and human trials employing therapeutic editing are already underway **[1]**. Genome editing by a CRISPR-associated endonuclease such as *Streptococcus pyogenes* Cas9 relies on the targeted induction of DNA double strand breaks (DSBs), leading to the recruitment of DNA repair factors that repair and potentially modify the genome **[2]**. However, unintended, off-target DNA damage and mutagenesis remain leading concerns for safety and applicability. Therefore, accurate and sensitive methods for discovery of CRISPR off-target activity are essential, especially for in vivo and therapeutic applications **[3]**.

There are numerous methods for detecting off-target CRISPR activity, but the majority are limited to purified DNA **[4-7]** or restricted cellular systems such as immortalized cell lines or reporter cells **[8-10]**. Measurements in these systems may not translate to in vivo and clinical applications. For example, Cas9 behavior such as binding kinetics is very different in vitro **[11]**, and the epigenome, which is highly divergent between different cell types **[12]**, strongly influences CRISPR genome editing activity **[13]**. Therefore, off-target discovery directly in clinically translatable ex vivo and in vivo model systems is highly desired, but the few methods compatible with these systems are constrained by limited sensitivity **[14-16]**.

A prior method for off-target detection in vivo, DISCOVER-Seq **[14]**, works by detecting the genome-wide localization of MRE11, a DNA repair factor recruited to Cas9 DSB sites, using chromatin immunoprecipitation sequencing (ChIP-seq) **[17,18]**. However, sensitivity is relatively low, likely because Cas9 editing is not synchronized and MRE11 resides on DNA only transiently during active repair **(Fig. 1a)**. We hypothesized that if DNA repair could be pharmacologically modulated to encourage MRE11 residence, then MRE11 would accumulate at every Cas9-targeted site in all cells, thus enhancing detection sensitivity with ChIP-seq **(Fig. 1b)**.

**Figure 1:**
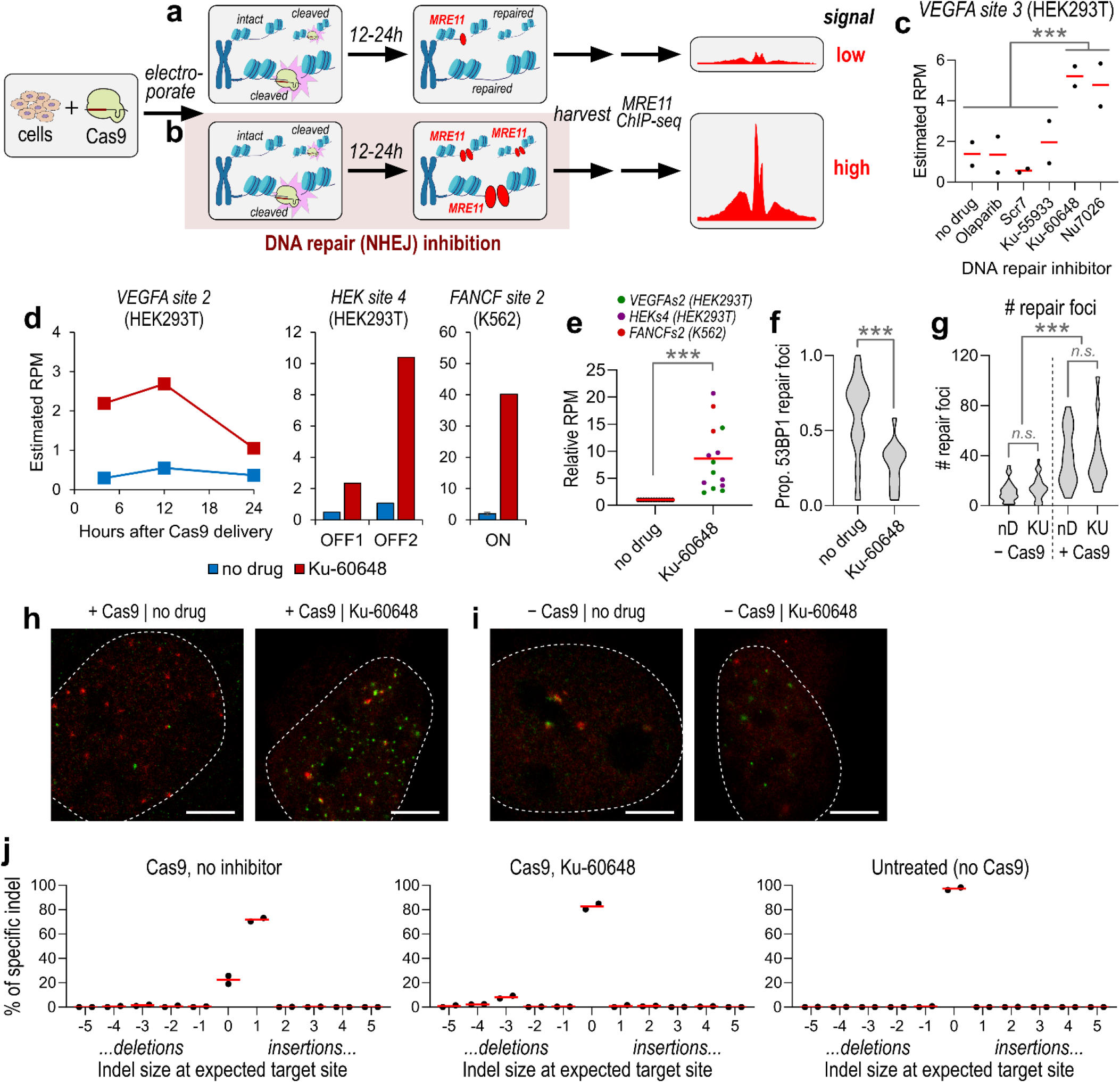
Effect of DNA-PKcs inhibition on DNA repair at CRISPR targeted locations. **a-b**, Schematic of genome-wide CRISPR off-target detection using MRE11 ChIP-seq. (a) Cells are unsynchronized, so only some cells have MRE11 at Cas9 cut sites at a given time (DISCOVER-Seq). (b) Inhibition of NHEJ directs DNA repair to slower, MRE11-dependent pathways. **c**, Effect of repair factor inhibition on MRE11 residence at the *VEGFA site 3* on-target site, measured by ChIP-qPCR estimating “reads per million” (RPM) enrichment at 12 hours after Cas9 delivery in HEK293T cells. ‘no drug’ means no repair factor inhibition, ‘Olaparib’ is a poly ADP ribose polymerase (PARP) inhibitor, ‘Scr7’ is a DNA Ligase IV inhibitor, ‘Ku-55933’ is an Ataxia–telangiectasia mutated (ATM) serine/threonine kinase inhibitor, and ‘Ku-60648’ and ‘Nu7026’ are DNA-dependent protein kinase catalytic subunit (DNA-PKcs) inhibitors. Each point corresponds to a different biological replicate of a sample exposed to the DNA repair inhibitor listed in the x-axis. Red line is mean of two biological replicates. Samples with DNA-PKcs inhibition (n = 4) have significantly higher estimated RPM compared to samples without DNA-PKcs inhibition (n = 8) using two-sided Student’s t-test (p < 1E-4). **d**, Increased MRE11 residence upon DNA-PKcs inhibition using Ku-60648 (red) versus without inhibition (blue), measured by ChIP-qPCR. Measured (left plot) over multiple time points (4 h, 12 h, 24 h) after delivery of Cas9 targeting *VEGFA site 2* in HEK293T, (middle plot) with Cas9 targeting *FANCF site 2* in K562 at 12 h, and (right plot) with Cas9 targeting *HEK site 4* in HEK293T at 12 h. Plots display the mean over two biological replicates for left and middle plots, and one replicate for the right plot. **e**, Plot of estimated RPM enrichment normalized to the no drug sample from data in panel d, for sample pairs with (‘Ku-60648’) or without (‘no drug’) DNA-PKcs inhibition. Normalized RPM enrichments with DNA-PKcs inhibition was significantly higher than without inhibitor (p = 0.0001), using Wilcoxon matched-pairs signed rank test, Red line indicates mean of n = 14 total samples pooled from panel d; green points are HEK293T, *VEGFA site 2*; purple points are HEK293T, *HEK site 4*; red points are K562, *FANCF site 2*. **f**, The proportion of 53BP1 foci relative to BRCA1 as detected by STED in cells exposed to Cas9 targeting a multi-target gRNA with 126 genome-wide target sites. N = 98 cells from 4 biological replicates, p < 1E-4 using two-sided Wilcoxon rank sum test. **g**, The number of repair foci (53BP1 or BRCA1) as detected by STED in cells with or without Cas9 (‘+ Cas9’ or ‘– Cas9’, respectively), with or without Ku-60648 (‘KU’ vs ‘nD’, respectively), targeting 126 genome-wide sites with a multi-target gRNA. N = 98 cells from 4 biological replicates, p < 1E-4 using two-sided Wilcoxon rank sum test between ‘– Cas9’ and ‘+ Cas9’. Difference in # of foci in each group was not significant (left, p = 0.15; right, p = 0.95). **h-i**, Representative images for panel g. Red labels 53BP1, green labels BRCA1. Scale bar corresponds to 5 μm. **j**, Histogram of insertion-deletion mutations (indels) or no mutations (‘0’) at 48 hours after Cas9 editing of *ACTB*, either without DNA-PKcs inhibitor (‘Cas9, no inhibitor’) or with inhibitor (‘Cas9, Ku-60648’). Untreated cells not exposed to Cas9 is shown for reference (‘Untreated (no Cas9)’).

In this study, we discovered that a small molecule inhibitor of DNA repair **[19]** greatly increases MRE11 residence on genomic DNA, enhancing the sensitivity of CRISPR off-target detection using MRE11 ChIP-seq. We demonstrated improved discovery of CRISPR-targeted sites in numerous contexts, including in immortalized cell lines, primary human cells, and mice at clinically relevant targets. Together, we developed the most sensitive method to-date for CRISPR off-target detection that is directly suitable for translational applications **[20]**.

## RESULTS

### Effects of DNA-PKcs inhibition on the CRISPR-mediated DNA damage response

To identify inhibitors of DNA repair **[20]** that can modulate MRE11 residence, we first delivered Cas9 with guide RNA (gRNA) targeting *VEGFA site 3* into HEK293T cells. *VEGFA site 3* and most other gRNAs used in this study were chosen because they have been well-validated in prior off-target detection methods **[8-10]**. We exposed cells to one of five DNA repair inhibitors, then performed ChIP with quantitative PCR (qPCR) after 12 hours to measure MRE11 recruitment at the target site. Inhibition of Poly (ADP-ribose) polymerase (PARP) and ATM serine/threonine kinase (ATM) with Olaparib and Ku-55933, respectively, did not exhibit a clear effect, whereas DNA Ligase IV inhibition with Scr7 suppressed MRE11 recruitment **(Fig. 1c) [21]**. Notably, blocking non-homologous end-joining (NHEJ) by inhibiting DNA-PKcs using Ku-60648 **[19,22]** or Nu7026 **[23]** significantly increased MRE11 recruitment at the target site (p < 1E-4; two-sided Student’s t-test) **(Fig. 1c)**. The effect of DNA-PKcs inhibition was consistent across multiple time points (4 hours, 12 hours, 24 hours), three other gRNAs (*VEGFA site 2, HEK site 4, FANCF site 2*), and/or another cell line (K562) (p < 0.001; two-sided Wilcoxon matched-pairs signed rank test) **(Fig. 1d-e)**. These results suggest that blocking NHEJ with DNA-PKcs inhibition greatly boosts MRE11 residence at Cas9-targeted sites.

We aimed to better characterize the effect of DNA-PKcs inhibition on repair of Cas9-mediated DNA damage. First, we used super-resolution stimulated emission depletion (STED) microscopy **[24,25]** to measure the localization of 53BP1 and BRCA1 foci after Cas9-induced DNA breaks in U2OS cells. 53BP1 corresponds to activation of the NHEJ pathway, whereas BRCA1 is implicated in MRE11-dependent homology directed repair (HDR) or microhomology-mediated end joining (MMEJ) **[26,27]**. Using a multi-target gRNA targeting over 100 locations **[28]**, DNA-PKcs inhibition using Ku-60648 led to a significant reduction in 53BP1 foci relative to BRCA1, consistent with suppression of NHEJ in favor of HDR/MMEJ (p < 1E-4; two-sided Wilcoxon rank sum test) **(Fig. 1f)**. Ku-60648 in the absence of Cas9 did not change the number of DNA damage (53BP1 and BRCA1) foci detectable by STED, suggesting that Ku-60648 alone does not induce DNA damage inside cells **(Fig. 1g-i)**. Additionally, we used a complementary assay to determine the type of insertion-deletion mutations (indels) by Sanger sequencing after 3 days of Cas9 targeting *ACTB* in HEK293T cells **[17]**. Exposure to Ku-60648 altered indel outcomes, from +1 insertions associated with NHEJ in favor of larger -3 deletions from MMEJ **(Fig. 1j)**. Together, these results confirm that DNA-PKcs inhibition with Ku-60648 blocks the NHEJ repair pathway in favor of MRE11-associated HDR and MMEJ pathways, therefore boosting MRE11 residence.

### MRE11 ChIP-seq with DNA-PKcs inhibition increases off-target detection sensitivity

Next, we determined whether increased MRE11 residence with DNA-PKcs inhibition can improve the sensitivity of CRISPR off-target discovery. At 12 hours after delivery of Cas9 with *FANCF site 2* gRNA into K562 cells, we performed ChIP-seq for MRE11 followed by the BLENDER bioinformatics pipeline **[14]** to detect all Cas9 target sites genome-wide. Sequencing samples with or without DNA-PKcs inhibition were always normalized to the same number of reads for appropriate comparison. Treatment with Ku-60648 significantly increased MRE11 ChIP-seq enrichment at all discovered on- and off-target sites (p < 1E-3; two-sided Wilcoxon rank sum test), as measured by the number of reads within a 1.5 kb region around the target site per million total reads, *i*.*e*., reads per million (RPM) **(Fig. 2a-d**). MRE11 ChIP-seq enrichment at a specific target site can be visualized as a histogram of base pair coverage along the genome; it exhibits two peaks on each side of the cut site because paired-end Illumina sequencing only reads the ends of DNA fragments that are enriched around the cut site **(Fig. S1a) [14]**. MRE11 levels 10 kb away from the target sites did not significantly increase (p ≥ 0.18; two-sided Wilcoxon rank sum test), further supporting the lack of additional DNA damage caused by the inhibitor itself **(Fig. 2e-f)**.

**Figure 2:**
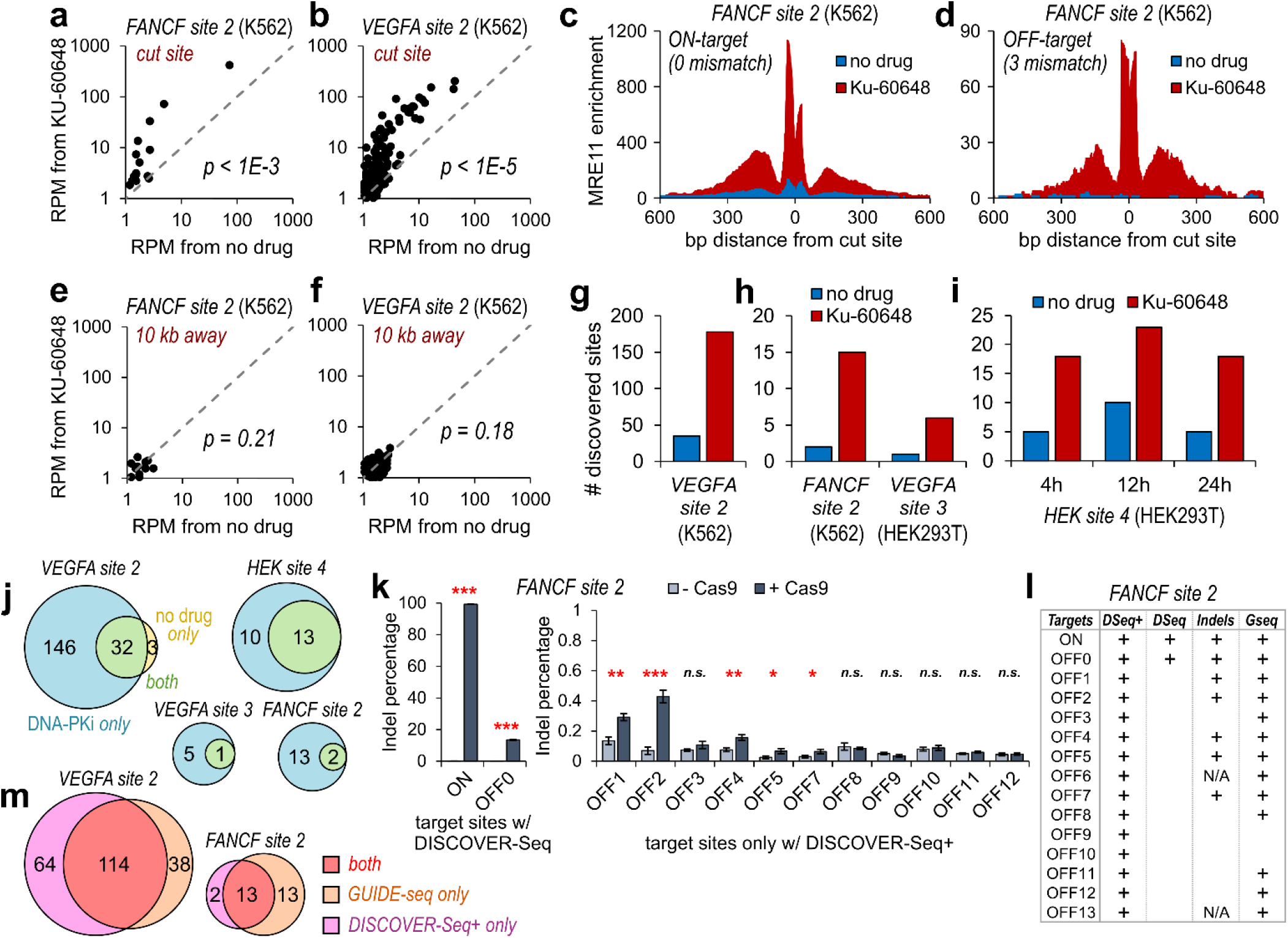
DNA-PKcs inhibition increases the sensitivity of CRISPR off-target detection. **a-b**, Plots of MRE11 ChIP-seq enrichment (number of reads within a 1.5 kb window centered at the cut site, per 1 million total reads, i.e., reads per million or RPM) for samples with (y-axis) or without (x-axis) DNA-PKcs inhibition at all (a) *FANCF site 2* or (b) *VEGFA site 2* Cas9 target sites in K562 detected from the DNA-PKcs inhibited samples. Each point in the plot (15 in panel a, 178 in panel b) corresponds to a putative target site. Significant differences (p < 1E-3 or p < 1E-5) between y-axis and x-axis values were determined using two-sided Wilcoxon rank sum test. **c-d**, Genome browser visualization of MRE11 enrichment at an (c) on-target and (d) representative off-target position from K562 with Cas9 targeting *FANCF site 2*, with (red) or without (blue) DNA-PKcs inhibition. **e-f**, Same as panels a and b, at positions 10 kb downstream from the actual cut site, to measure background enrichment adjacent to cut sites. MRE11 enrichment with (y-axis) versus without (x-axis) DNA-PKcs inhibition at the adjacent background locations was not significantly different (p = 0.21 or 0.18), determined using two-sided Wilcoxon rank sum test. **g**, Number of discovered off-target sites with (red) or without (blue) DNA-PKcs inhibition for *VEGFA site 2*. Quantification of Figure S1b. **h-i**, Number of discovered off-target sites with (red) or without (blue) DNA-PKcs inhibition for (h) *VEGFA site 3, FANCF site 2*, and (i) *HEK site 4* gRNAs. Quantification of Figure S1c. **j**, Venn diagram illustrating overlap in the identity of Cas9 target sites discovered from samples with DNA-PKcs inhibition (‘DNA-PKi only’; light blue), without DNA-PKcs inhibition (‘no drug only’; light yellow), or found in both samples (‘both’; light green). 4 gRNAs were evaluated. **k**, (left plot) Measurement of insertion-deletion mutations (indels) at the on-target and sole off-target site (OFF0) discovered by the original DISCOVER-Seq, for K562 cells with the *FANCF site 2* gRNA, with or without Cas9 (‘+ Cas9’ or ‘-Cas9’, respectively). (right plot) Measurement of indels at off-target sites exclusively discovered by DISCOVER-Seq+. Plots display the mean of 3 biological replicates, error bars represent ±1 standard deviation from mean. n.s. indicates not significant, * indicates p<0.05, ** indicates p<0.01, and *** indicates p<0.001 using Student’s t-test. **l**, For the 15 target sites (1 on-target, 14 off-targets) of the *FANCF site 2* gRNA identified by DISCOVER-Seq+ (‘DSeq+’), the chart shows which sites are also identified by DISCOVER-Seq alone (‘+’ under ‘DSeq’), which sites have indels detectable by targeted deep sequencing (‘+’ under ‘Indels’), and which sites were also detectable by GUIDE-seq (‘+’ under ‘Gseq’). Target sites labeled with ‘N/A’ under ‘Indels’ were unable to be successfully amplified by PCR for targeted sequencing. **m**, Venn diagram illustrating overlap in the identity of *VEGFA site 2* and *FANCF site 2* target sites identified by DISCOVER-Seq+ versus GUIDE-seq.

To reduce the likelihood of reporting false positive sites, MRE11 ChIP-seq was also performed on cells with the same experimental conditions except *without* Cas9. The final set of off-target sites is therefore determined as the set from the sample with Cas9 subtracted by the set from the corresponding sample without Cas9. For the *VEGFA site 2* gRNA, 178 sites were discovered with DNA-PKcs inhibition using Ku-60648, which is an over 5-fold increase compared to 35 sites discovered without DNA-PKcs inhibition (i.e., DISCOVER-Seq) **(Fig. 2g; Fig. S1b)**. Improved performance with Ku-60648 was consistent across different gRNAs and multiple time points **(Fig. 2h-i; Fig S1c)**. The discovered sites with Ku-60648 included almost all the sites identified using DISCOVER-Seq alone **(Fig. 2j)**. Reassuringly, only a small minority (average of 1.7%) of the initial sites were also found in corresponding negative control samples without Cas9, and therefore deemed to be false positives and removed **(Fig. S2a-b)**. We therefore use the term DISCOVER-Seq+ to denote CRISPR off-target discovery that combines MRE11 ChIP-seq (i.e., DISCOVER-Seq) **[14]** with DNA-PKcs inhibition to achieve improved detection sensitivity.

Next, we assessed if any of the new sites discovered by DISCOVER-Seq+ harbor evidence of mutagenesis after CRISPR genome editing. For the *FANCF site 2* gRNA, DISCOVER-Seq+ identified 15 target sites, compared to only two with DISCOVER-Seq **(Fig. 2j)**. We exposed cells to Cas9 targeting *FANCF site 2* for 4 days (without Ku-60648), then measured indel mutations at each discovered target site using deep amplicon sequencing **[29]**. Of the 13 off-target sites exclusively discovered by DISCOVER-Seq+, five exhibited detectable indels by amplicon sequencing **(Fig. 2k)**. These results demonstrate that DISCOVER-Seq+ identified new off-target sites with evidence of indel mutations, which DISCOVER-Seq alone failed to detect. Although some newly discovered off-target sites lacked detectable indel mutations by amplicon sequencing, they are still essential to identify because DSBs, even in the absence of mutagenesis, are detrimental to the cell **[21,28]**.

The validity of DISCOVER-Seq+ off-target sites was further confirmed by comparing to published results by an independent technique, GUIDE-seq **[8]**, which is notably not compatible with primary cells or in vivo applications **[14]**. For both the *FANCF site 2* and *VEGFA site 2* gRNAs, half or more of target sites found by DISCOVER-Seq+ were also found by GUIDE-seq, and vice versa. **(Fig. 2l-m)**. These results demonstrate robust overlap in the discovered target sites between GUIDE-seq and DISCOVER-Seq+, providing external validity for DISCOVER-Seq+.

### DISCOVER-Seq+ in editing of primary human cells

We further evaluated the utility of DISCOVER-Seq+ in three emerging clinical applications: patient-derived iPSC editing, generating engineered T cells for cancer immunotherapy, and in vivo characterizations of CRISPR-based therapies in mouse models. All three applications share a critical reliance on off-target identification to ensure safety with clinical translation. First, we used DISCOVER-Seq+ to improve off-target detection in induced pluripotent stem cells (iPSCs). DISCOVER-Seq+ in WTC-11 iPSCs **[30]** discovered over 2-fold more off-target sites at *VEGFA site 2* compared to DISCOVER-Seq **(Fig. 3a)**. At all discovered off-target sites, MRE11 ChIP-seq enrichment was also significantly increased (p < 1E-5; two-sided Wilcoxon rank sum test) **(Fig. 3b-c; Fig. S2c)**.

**Figure 3:**
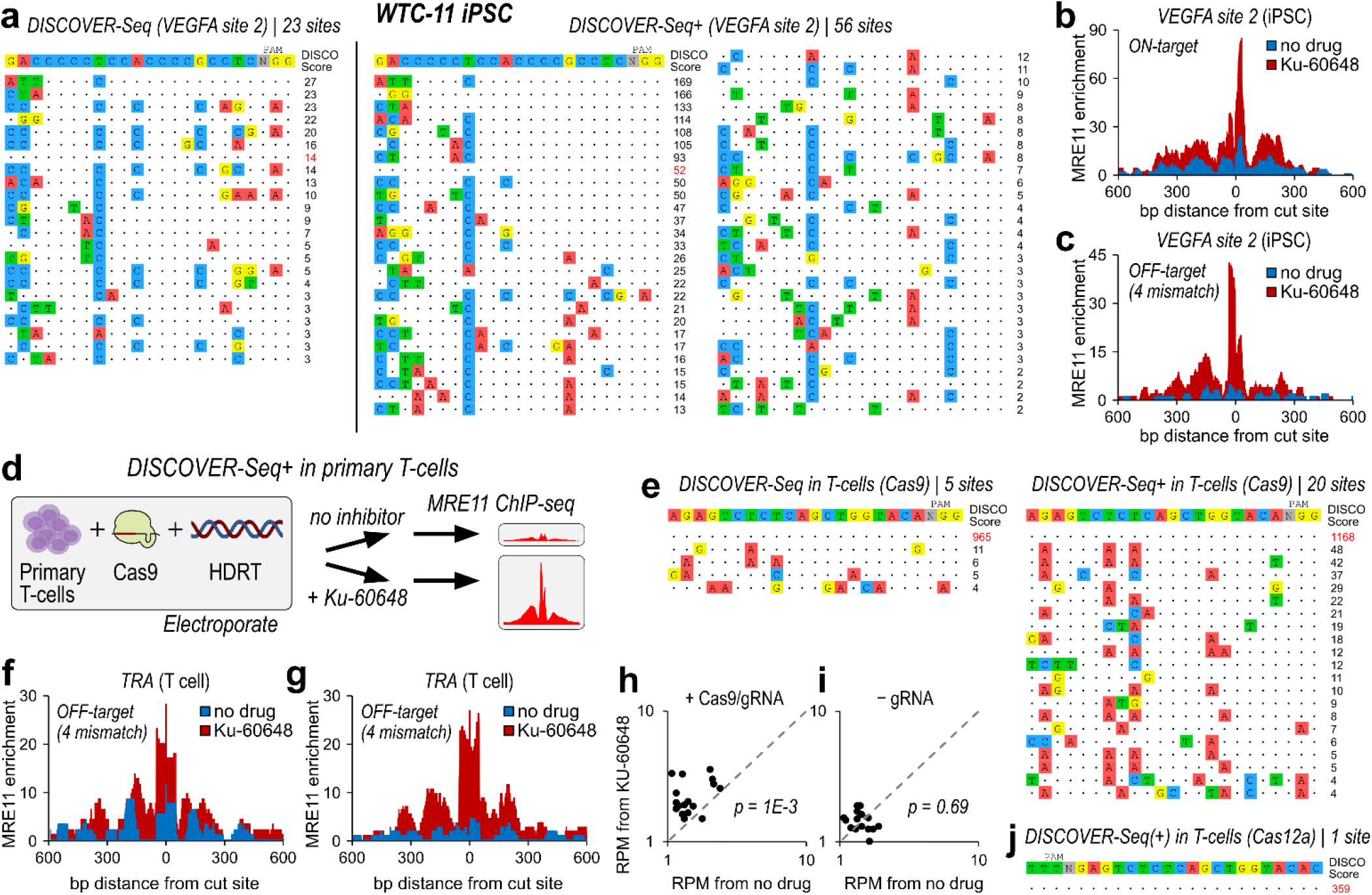
DISCOVER-Seq+ in human iPSCs and primary T cells. **a**, *VEGFA site 2* Cas9 target sites detected using DISCOVER-Seq (left) versus DISCOVER-Seq+ (right) in WTC-11 iPSCs. **b-c**, Genome browser visualization of MRE11 enrichment at an (b) on-target and (c) representative off-target position with 4 mismatches (‘4 mm’) in WTC-11 iPSCs with Cas9 targeting *VEGFA site 2*. DISCOVER-Seq+ data in red (with Ku-60648), DISCOVER-Seq data in blue (with no drug exposure). **d**, Schematic of the DISCOVER-Seq+ protocol in the knock-in of a cancer neoantigen-specific, transgenic TCR (tgTCR) into the *TRA* locus of primary human T cells. **e**, *TRA* Cas9 target sites in primary T cells detected using DISCOVER-Seq (left) versus DISCOVER-Seq+ (right). **f-g**, Genome browser visualization of MRE11 enrichment at two representative 4-mismatch (‘4mm’) off-target positions in primary human T cells with Cas9 targeting *TRA* for knock-in of a tgTCR template. DISCOVER-Seq+ data in red (with Ku-60648), DISCOVER-Seq data in blue (with no drug exposure). **h-i**, (h) Plots of MRE11 ChIP-seq reads-per-million enrichment within a 1.5 kb window for samples with (y-axis) or without (x-axis) DNA-PKcs inhibition, at all *TRA* Cas9 off-target sites in primary human T cells from the DNA-PKcs inhibited samples. Each point in the plot (20 total) corresponds to a putative target site. Significant differences (p < 1E-3 or p < 1E-5) between y-axis and x-axis values were determined using two-sided Wilcoxon rank sum test. (i) Same as panel h, for cells delivered with Cas9 but without gRNA (negative control). **j**, *TRA* Cas12a (Cpf1) target sites in primary T cells. The results were the same between DISCOVER-Seq and DISCOVER-Seq+; only the on-target site was detected.

Next, we applied DISCOVER-Seq+ ex vivo to knock-in of a cancer neoantigen-specific transgenic T-cell receptor (tgTCR) construct into primary human T cells **[31-32]**. We electroporated Cas9 targeting *TRA* (T Cell Receptor Alpha Locus) along with a 4699 bp homology-directed repair template (HDRT) encoding a tgTCR specific for HLA-A*02 loaded with mutant p53 R175H peptide **[31]**, then performed DISCOVER-Seq+ 12 hours later **(Fig. 3d)**. The specific R175H mutation that is targeted by the tgTCR is the most prevalent p53 gain-of-function mutation in human cancers **[33]**. DISCOVER-Seq+ (with Ku-60648) identified 20 off-target sites genome-wide compared to 4 with DISCOVER-Seq **(Fig 3e)**, and led to significantly greater MRE11 enrichment at all discovered sites (p = 1E-3; two-sided Wilcoxon rank sum test) **(Fig. 3f-h)**. In contrast, samples without Cas9 exhibited no change in enrichment with Ku-60648, further confirming that the inhibitor alone does not induce damage (p = 0.69; two-sided Wilcoxon rank sum test) **(Fig. 3i)**.

DISCOVER-Seq+ also has the potential to compare off-target profiles between different types of CRISPR nucleases. As a proof of concept, we compared the performance of Cas9 to Cas12a (Cpf1), targeting the same position in *TRA*. DISCOVER-Seq+ at 12 hours after Cas12a editing only identified the on-target site and no off-target sites **(Fig. 3j)**, consistent with the improved specificity of Cas12a **[34]**. Flow cytometry for tgTCR expression after 7 days in T cells without Ku-60648 exposure showed similar tgTCR integration efficiencies of 8.4% for Cas12a and 9.7% for Cas9 **(Fig. S2d-f)**. Together, our preliminary analysis using DISCOVER-Seq+ revealed that Cas12a maintained adequate tgTCR integration rates while eliminating detectable off-target damage. Importantly, these experiments demonstrated that DISCOVER-Seq+ is directly compatible with CRISPR knock-in using a homology template in primary human T cells.

### DISCOVER-Seq+ in vivo

Finally, we evaluated DISCOVER-Seq+ in vivo by targeting the cardiovascular risk gene *PCSK9* in mouse liver **[35]**. We retro-orbitally injected adenovirus encoding Cas9 and *PCSK9* gRNA into C57BL/6J mice, followed by peritoneal injection of either 25 mg/kg Ku-60648 (i.e., DISCOVER-Seq+) or vehicle (i.e., DISCOVER-Seq) twice daily (b.i.d.) **(Fig. 4a)**. We selected the specific *PCSK9* gRNA that was also used in the original DISCOVER-Seq study for direct comparison **[13]**. Ku-60648 has been evaluated as a drug for chemo-sensitization in cancer therapy, exhibits good pharmacokinetics, and strongly penetrates tissue including tumors **[22]**. Mice were sacrificed after 24 hours to harvest the liver for MRE11 ChIP-seq. DISCOVER-Seq+ mice exhibited increased MRE11 ChIP-seq signal in their liver compared to those without DNA-PKcs inhibition (p < 1E-4; two-sided Wilcoxon rank sum test) **(Fig. 4b-d; Fig. S2g)**. An average of 27 target sites were identified with DISCOVER-Seq+ compared to 18 sites with DISCOVER-Seq across 5 biological replicates (p < 0.01; two-sided Student’s t-test) **(Fig. 4e)**. The identified sites strongly overlap between the two methodologies and with sites identified in the original DISCOVER-Seq study **(Fig. 4f) [14]**. Pooling sequencing reads across all 5 replicates identified 98 target sites with DISCOVER-Seq+ versus 49 with DISCOVER-Seq **(Fig. 4g)**. Together, these results demonstrate that DISCOVER-Seq+ is compatible with direct measurement of genome-wide off-target editing in vivo.

**Figure 4:**
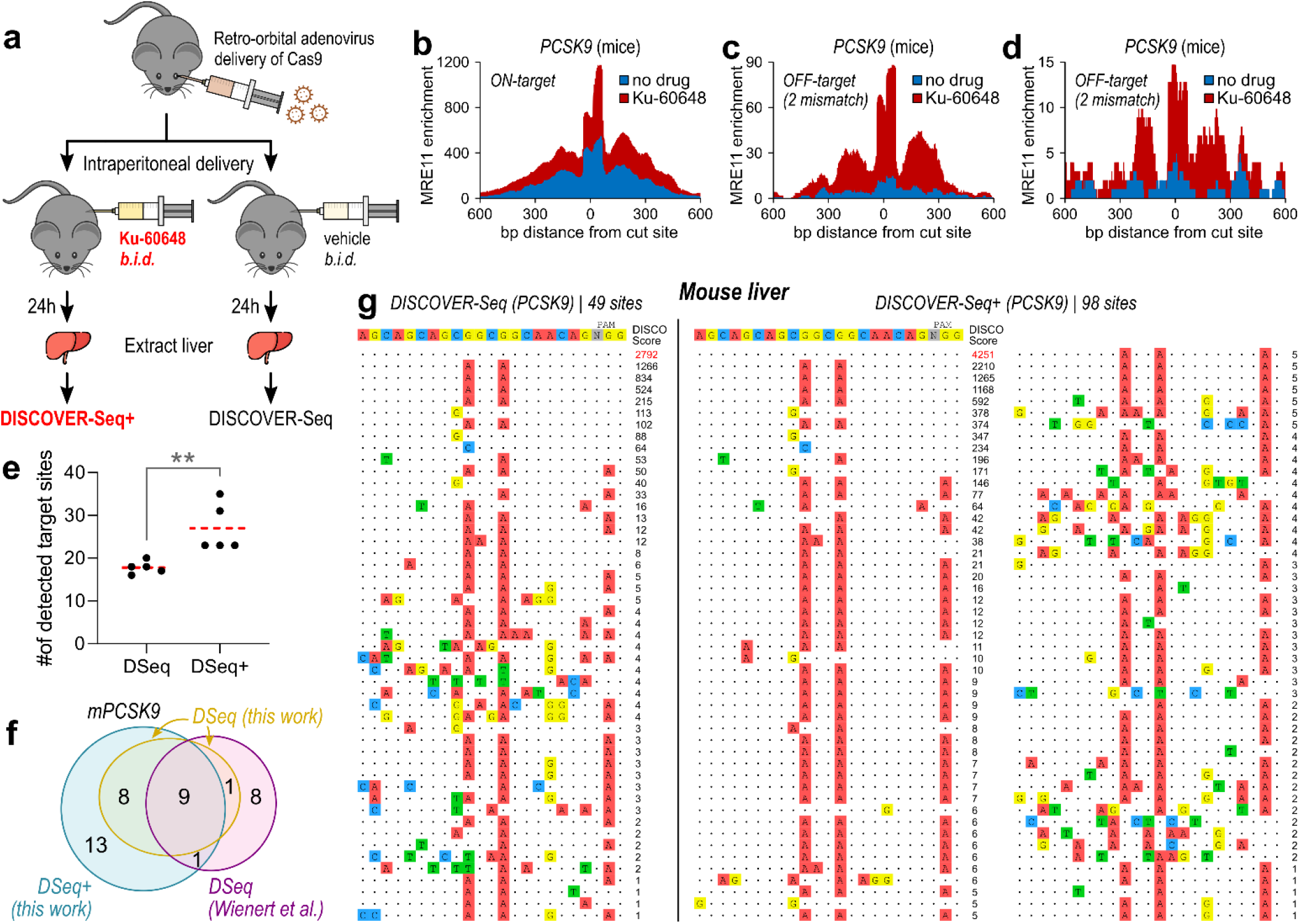
DISCOVER-Seq+ in mice. **a**, Schematic of DISCOVER-Seq+ protocol in mice. **b-d**, Genome browser visualization of MRE11 enrichment at the (b) *PCSK9* on-target site (‘ON-target’) and (c,d) two off-target sites (‘OFF-target’) with 2 mismatches each (‘2 mm’) in the liver of mice transduced with adenovirus expressing Cas9 targeting *PCSK9*. Mice were dosed twice a day (*b*.*i*.*d*.) with either 25 mg/kg Ku-60648 (‘Ku-60648’; red) or with vehicle (‘no drug’; blue). **e**, Number of detected genome-wide target sites in the mouse genome mm10 with Cas9/gRNA targeting *PCSK9*, identified using DISCOVER-Seq (DSeq) versus DISCOVER-Seq+ (DSeq+). N=5 biological replicates (mice) were used for each condition; DISCOVER-Seq+ detected significantly more target sites than DISCOVER-Seq, using two-sided Student’s t-test (p = 0.0079). **f**, Venn diagram illustrating overlap in the mouse *PCSK9* target sites identified by in vivo DISCOVER-Seq+ in this work *– ‘DSeq+ (this work)’* (blue), in vivo DISCOVER-Seq in this work *– ‘DSeq (this work)’* (yellow), and the original DISCOVER-Seq manuscript *– ‘DSeq (Wienert et al*.*)’* (purple). All the target sites identified by *‘DSeq (this work)’* are also found in the other two groups. **g**, *PCSK9* Cas9 target sites detected using DISCOVER-Seq (left) versus DISCOVER-Seq+ (right) in mouse liver. Off-target detection for each condition was performed on sequencing data pooled across 5 biological mouse replicates.

## DISCUSSION

This study designed and validated DISCOVER-Seq+, the most sensitive method to-date for detecting CRISPR off-target activity in primary cells and in vivo. As CRISPR becomes an increasingly feasible approach for therapeutic genome editing **[1,20,31,32,35-37]**, evaluation of CRISPR off-target activity directly in clinically translatable applications is crucial. Even if a specific CRISPR gRNA does not have detectable off-target activity in one particular cell type, this may not translate to other cell types, tissues, or organisms. For example, despite using the same *VEGFA site 2* gRNA, there are differences in target sites between 3 different cell lines (HEK293T, K562, and WTC-11 iPSC) **(Fig. S2h-i)**. Directly measuring off-target sites in the system of interest, whether in primary T cells, mice, or even non-human primates, is therefore essential.

DISCOVER-Seq+ may find use in numerous clinical applications. For in vivo liver editing, such as to prevent cardiovascular disease **[35]** or to treat Transthyretin (ATTR) Amyloidosis **[36]** in a particular patient, initial off-target validation of gRNAs in non-human primates along with personalized validation in primary liver cells derived from that individual patient may be essential before therapy. Ex vivo generation of engineered T cells **[31,32,37]** further highlights a pressing need for personalized evaluation of off-target sites; since each patient receives a bespoke T cell line derived from their own cells, a precision medicine approach will be required to ensure each patient’s off-target profile is acceptable before therapy.

Compared to DISCOVER-Seq+, other methods for off-target detection are limited to in vitro measurements (e.g., CIRCLE-Seq, Digenome-seq) **[4-7]**, limited to immortalized cell lines (e.g., GUIDE-seq) **[8-10]**, or exhibit limited sensitivity (e.g., original DISCOVER-Seq) **[14]**. Therefore, by combining unparalleled detection sensitivity with high versatility in ex vivo and in vivo applications, we believe DISCOVER-Seq+ will find widespread use as the state-of-the-art technology for off-target detection in translational and clinical applications of therapeutic CRISPR editing.

We extensively evaluated the possibility of false positive detection with DISCOVER-Seq+. Theoretically, false positives could arise from non-Cas9 mediated DNA damage and/or from the ChIP-seq readout itself. These false positive sites are experimentally determined using MRE11 ChIP-seq in matching samples without Cas9, and removed from consideration. Importantly, we found the percent of false positives to be very small at approximately 1.7% across all evaluated gRNAs **(Fig. S2a-b)**, which is reassuring. To further exclude the likelihood of false positives in our protocol, we performed numerous experiments that showed the DNA-PKcs inhibitor is unlikely to induce DNA damage by itself: (1) cells with Ku-60648 in the absence of Cas9 gRNA had no change in the number of DNA damage foci as measured by super-resolution STED microscopy **(Fig. 1f-i)**, or in the level of MRE11 recruitment as measured by ChIP-seq **(Fig. 3i)**, (2) adding Ku-60648 to cells with Cas9 did not increase DNA damage in regions outside of predicted off-target sites **(Fig. 2e-f)**, and (3) off-target sites found by DISCOVER-Seq+ but not by DISCOVER-seq were also independently found using alternative methods such as GUIDE-seq **(Fig. 2m)**.

DISCOVER-Seq+ identified off-target sites with evidence of mutagenesis that DISCOVER-Seq alone failed to detect **[8,38]**. Furthermore, because DISCOVER-Seq+ directly measures off-target *DNA damage* rather than *mutagenesis*, it also discovered sites that did not have detectable indel mutations **(Fig. 2k-l)**. This is important for three reasons: (1) One study found that less than 15% of DSBs convert to indels in a single damage cycle **[17]**; therefore, off-target sites that are not frequently damaged may not result in indels that can appear in amplicon sequencing-based queries. (2) DNA damage such as DSBs are highly detrimental regardless of mutagenesis, especially to primary cells, by inducing widespread epigenetic changes and perturbing native cellular functions such as cell division and transcription **[21,28]**. (3) Evaluating indels alone may also miss complex DNA damage outcomes such as large deletions and translocations **[8,39]**. Furthermore, deep amplicon sequencing may miss rarer off-target sites due to a lack of enrichment of altered DNA molecules and insufficient sequencing depth **[40]**. For all these reasons, enrichment for sites with evidence of CRISPR-induced DNA damage is a more holistic readout of off-target activity than indels alone. In summary, DISCOVER-Seq+ directly detects genome-wide CRISPR-induced DNA damage, regardless of whether these sites become mutated, with unprecedented sensitivity and versatility.

DNA-PKcs inhibition could also influence the performance of other methods to detect CRISPR activity. BLISS/BLESS **[9]** could be improved with DNA-PKcs inhibition by increasing the quantity of unrepaired DSBs at the time of evaluation. In contrast, GUIDE-seq **[8]** would likely be impaired because incorporation of its double-stranded oligodeoxynucleotide (dsODN) relies on NHEJ, which is directly inhibited by DNA-PKcs inhibitors. Future studies should explore whether other strategies of inhibiting DNA repair may improve detection of CRISPR off-target activity.

Limitations of DISCOVER-Seq+ include the need for the DNA-PKcs inhibitor to exert its effect, including in vivo. We believe this is not a major concern because Ku-60648 is a small molecule previously shown to have good bioavailability and tissue penetration, including into tumors **[22,23]**. In contrast, the main barrier to applicability in other organs in vivo remains the efficiency of CRISPR delivery to non-liver organs **[1,14,35]**. In addition, the improvement in the number of discovered off-target sites with DNA-PKcs inhibitor in mice **[14]** is a factor of two, which is lower than in cell lines. This may be due to reduced inhibitor or Cas9 concentrations in the liver; further optimization of drug dosing and Cas9 delivery may improve performance.

Identifying off-target genome editing in therapeutically relevant systems is a major barrier to clinical translation of CRISPR systems. By leveraging MRE11 ChIP-seq with DNA-PKcs inhibition, DISCOVER-Seq+ provides the highest detection sensitivity to-date in systems ranging from ex vivo editing of primary human cells to in vivo editing of mice, setting the standard for genome-wide CRISPR off-target discovery **(Table S1)**. DISCOVER-Seq+ has the potential to validate the specificity profile of genome editing at numerous stages of the therapeutic development pipeline, from cell lines and primary cells to mice and potentially non-human primates **[1,18,31,32,35-37]**.

## Acknowledgements

We thank members of the Taekjip Ha, Sua Myong, and Mark E. Anderson (M.E.A.) labs for support, in particular, Jonathan Granger and Elizabeth Luczak. This work was supported by grants from the National Institutes of Health (R35 GM 122569 and U01 DK 127432 to T.H.; T32 GM 136577 and F30 CA 254160 to R.S.Z.; T32 GM 136577 to B.J.M.; R35 HL 140034 to M.E.A.) and the National Science Foundation (PHY 1430124 to T.H.). Y.L. is supported by Research Corporation for Science Advance (RCSA) as a Cottrell Postdoctoral Fellow. M.F.K. is supported by the Jerome Greene Foundation and Cupid Foundation. A.M.G. is a Howard Hughes Medical Institute Awardee of the Life Sciences Research Foundation. T.H. is an investigator of the Howard Hughes Medical Institute.

## Author Contributions

R.S.Z. and T.H. conceived the study and oversaw experimental design. Y.L. and R.S.Z. identified the potent inhibitors and validated DISCOVER-Seq+ in immortalized cell lines and iPSCs. O.E.R.G. and R.S.Z. validated DISCOVER-Seq+ in mice. M.F.K., R.S.Z., and B.J.M. validated DISCOVER-Seq+ in primary T cells. R.S.Z., Y.L., and A.M-G. performed ChIP-seq. Y.L. performed immunofluorescence. L.L.S. performed deep amplicon sequencing. F.A-Z. performed structured illumination microscopy. R.S.Z. performed bioinformatics, data analysis, and wrote the manuscript with contributions from all authors. T.H. supervised the study.

## Competing interests

Johns Hopkins University has submitted a patent application on the method for off-target detection used in this study.

## Data availability

All sequencing data are uploaded to Sequence Read Archive under BioProject accession PRJNA801688. All other data are available within the paper and its supplementary information files.

## Code availability

Complete analysis code available on GitHub (https://github.com/rogerzou/DSeqPlus).

## METHODS

### Cell culture

HEK293T cells (ATCC^®^ CRL-3216™) and K562 cells (ATCC^®^ CCL-243™) were cultured at 37°C under 5% CO_2_ in Dulbecco’s Modified Eagle’s Medium (DMEM, Corning) supplemented with 10% FBS (Clontech), 100 units/mL penicillin, and 100 μg/mL streptomycin (DMEM complete). Cells were tested every month for mycoplasma.

A human induced pluripotent stem cell (hiPSC), the WTC-11 cell line (Kreitzer et al., 2013), was used for all iPS cell experiments in this study. We followed the guidelines of Johns Hopkins Medical Institute for the use of this hiPSC line. Briefly, frozen WTC11 cells were first thawed in 37°C water bath and washed in Essential 8 Medium (E8; Thermo Fisher Scientific, #A1517001) by centrifugation. After resuspension, WTC-11 cells were plated onto a 6 cm cell culture dish pre-coated with human embryonic cell (hES cell)-qualified matrigel (1:100 dilution, Corning #354277). Plate coating should be performed for at least 2 hours. Subsequently, 10 μM ROCK inhibitor (Y-27632; STEMCELL, #72308) was supplemented into the E8 medium to promote cell growth and survival. For subculture, WTC-11 cells were dissociated from the plate using Accutase (Sigma, #A6964) and passaged every 2 days. WTC-11 cells were maintained in an incubator at 37°C with 5% CO_2_.

### Mouse husbandry

All mouse studies were carried out in accordance with guidelines and approval of the Johns Hopkins University Animal Care and Use Committee (Protocol #MO20M274). Male C57BL/6J mice (The Jackson Laboratory, ME, USA) were housed in a facility with 12-hour light/12-hour dark cycle at 22 ± 1 °C and 40 ± 10% humidity. Teklad global 18% protein rodent diet and tap water were provided ad libitum.

### Immunofluorescence and imaging by STED microscopy

U2OS cells stably expressing Cas9-EGFP cells were seeded onto 35-mm, glass bottom dishes and transfected with mtgRNAs for 12-24 hrs. Cleavage was activated by UV light for 1 minute. To fix cells, 4 % of pre-warmed paraformaldehyde in 1X PBS was used for 10 min. After rinsing 3 times with 1X PBS, cell membrane permeabilization was done with Triton-X used for 10 min. Then, cells 2 % w/v BSA in 1x PBS was used for blocking for 1 hr and at room temperature. The primary antibodies, mouse anti-BRCA1 (D-9 sc-6954 Santa Cruz) and anti-53bp1 (Abcam, ab172580), were diluted (1:500) in 1x PBS and directly added into the imaging dish. After 1 hr incubation, the primary antibody was removed, and the sample was washed with 1x PBS three times. The samples were then incubated for 30 minutes with the secondary antibodies Alexa-594 and Atto-647N diluted (1:1000) in 1x PBS. Finally, the sample was rinsed three times and mounted with Prolong Diamond mounting media (Thermo Fisher Scientific) overnight.

All STED images were obtained using a home-built two-color STED microscope (Han and Ha, 2015; Ma and Ha, 2019). In short, a femtosecond laser beam with a repetition rate of 80 MHz from a Ti:Sapphire laser head (Mai Tai HP, Spectra-Physics) is split into two parts: one part produces an excitation beam coupled into a photonic crystal fiber (Newport) for wide-spectrum light generation. The beam is further filtered by a frequency-tunable acoustic optical filter (AA Opto-Electronic) for multi-color excitation. The other part of the laser pulse is temporally stretched to ∼300 ps (with two 15-cm-long glass rods and a 100-m long polarization-maintaining single-mode fiber, OZ optics), collimated and expanded, and wave-front modulated with a vortex phase plate (VPP-1, RPC photonics). This modulation produces a hollow STED spot generation to de-excite the fluorophores at the periphery of the excitation focus, thus improving the lateral resolution. The STED beam is set at 765 nm with a power of 120 mW at the back focal plane of the objective (NA=1.4 HCX PL APO 100×, Leica), and the excitation wavelengths are set as 594 nm and 650 nm for imaging Alexa-594 and Atto-647N labeled targets, respectively. Two avalanche photodiodes detect the fluorescent photons (SPCM-AQR-14-FC, Perkin Elmer). The images are obtained by scanning a piezo-controlled stage (Max311D, Thorlabs) controlled with the Imspector data acquisition program.

### Electroporation of Cas9 RNP and DNA-PKcs inhibitor delivery into cell lines and iPSCs

crRNA and tracrRNA sequences are in Table S2. 2 μL of 100 μM crRNA was mixed with 2 μL of 100 μM tracrRNA (Integrated DNA Technologies) and heated to 95°C for 5 min in a thermocycler, then allowed to cool on benchtop for 5 min. To form the RNP complex, 3 μL of 10 μg/μL (∼66 μM) of purified Cas9 was mixed with the annealed 4 μL 50 μM cr:tracrRNA, then 8 μL of dialysis buffer (20 mM HEPES pH 7.5, and 500 mM KCl, 20% glycerol) was mixed in for a total of 15 μL. This solution as incubated for 20 min at room temperature to allow for RNP formation.

HEK293T cells were maintained to a confluency of ∼90% prior to electroporation. 12 million cells were trypsinized with 5 min incubation in the incubator, then 1:1 of DMEM complete was added to inactivate trypsin. This mixture was centrifuged (3 min, 200 *× g*), supernatant removed, followed by resuspension of the cell pellet in 1 mL PBS, centrifugation (3 min, 200 *× g*), and finally complete removal of supernatant. 90 μL of nucleofection solution (16.2 μL of Supplement solution mixed with 73.8 μL of SF solution from SF Cell Line 4D-Nucleofector™ X Kit L) (Lonza) was mixed thoroughly with the cell pellet. The 15 μL RNP solution was mixed in along with 2 μL of Cas9 Electroporation Enhancer (Integrated DNA Technologies). The entirety of the final solution (approximately 125 μL) was transferred to one well of a provided cuvette rated for 100 μL. Electroporation was then performed according to the manufacturer’s instructions on the 4D-Nucleofector™ Core Unit (Lonza) using code CA-189. Some white residue may appear in the cell mixture after electroporation, but that is completely normal. A total of 400 μL of DMEM complete was used to completely transfer the cells out of the cuvette, before plating to culture wells pre-coated with 1:100 collagen. A minimum of 4 million cells are used for each ChIP. For time-resolved experiments, this means one electroporation equates to 3 samples.

For WTC-11 iPSCs, cells were dissociated from the plate using accutase (Sigma, #A6964). Electroporation was performed using the Lonza P3 Primary Cell 4D-Nucleofector™ X Kit L using code CA-137, on 10 million cells, and using 65 μL of the P3 solution mixture with EP enhancer per electroporation cuvette (compared to 90 μL of comparable SF solution mixture for HEK293T cells). After electroporation, cells were resuspended in E8 medium supplemented with 10 μM ROCK inhibitor (Y-27632; STEMCELL, #72308), and plated onto a 10 cm cell culture dish pre-coated with human embryonic cell (hES cell)-qualified matrigel (1:100 dilution, Corning #354277) for at least 2 hours.

To expose cells to DNA repair inhibitors, they were added to the culture media at a final concentration of 1 μM KU-60648 (1:2500 of 2.5 mM KU-60648), 20 μM Nu7026 (1:500 of 10 mM Nu7026), 10 μM Ku-55933 (1:10000 of 100 mM KU-55933), 1 μM Scr7 (1:10000 of 10 mM Scr7), or 10 μM Olaparib (1:1000 of 10 mM Olaparib). All stock solutions of drug were diluted in DMSO.

### Adenovirus and DNA-PKcs inhibitor delivery into mice

For *in vivo* gene delivery, 8-10 week old mice were anesthetized with 2.5% isofluorane/oxygen mixture. Mice received a single retro-orbital injection of 1 × 10^9^ infectious adenoviral particles (Ad-Cas9-U6-mPCSK9-sgRNA) in 100 μL sterile saline. Immediately following, mice received intraperitoneal delivery of KU-60648 dosed at 25 mg/kg (or vehicle only) in 100 μL of citrate buffer, or 100 μL of citrate buffer vehicle. Mice received a dose of KU-60648 or vehicle every 12 hrs via intraperitoneal injection.

### Extraction of mouse liver into cell suspension

At the experimental endpoint of 12 h, mice were anesthetized with isofluorane and euthanized via cervical dislocation. Liver tissue was harvested, washed 3x in 2 mL PBS with 1x protease inhibitor (Halt™ Protease Inhibitor Cocktail, Thermo), then disrupted the tissue in 1 mL PBS with 1x protease inhibitor using a loose-fitting Dounce homogenizer. For MRE11 ChIP-seq, homogenized tissue was placed on ice and used immediately.

### DISCOVER-Seq+/DISCOVER-Seq/MRE11 ChIP-seq

The protocol was adapted from previous literature (Wienert et al., 2019) and describes the reagents for one MRE11 ChIP-seq experiment.

For adherent cells, approximately 10 million cells were gently rinsed with room temperature PBS, washed off the plate using 10 mL DMEM with assistance from pipette squirts and cell scraper, then transferred to a 15 mL Falcon tube. For suspension cells, approximately 10 million cells were transferred to a 15 mL Falcon tube, spun down 200 *× g* for 1 min, decanted, then resuspended with 10 mL DMEM. 721 μL of 16% formaldehyde (methanol-free) was added and the tube was mixed by inversion in room temperature – 7 min for WTC-11 iPSCs, 12 min for HEK293T cells, or 15 min for K562 cells. For mouse liver, 300 μL of Dounce homogenized mouse liver was diluted into 10 mL PBS. 721 μL of 16% formaldehyde (methanol-free) was added and mixed by inversion in room temperature for 10 min.

Afterwards, 750 μL of 2 M glycine was added to quench the formaldehyde. Cells were spun down with 1,200 *× g* at 4°C for 3 min, then washed with ice-cold PBS twice, spinning down with the same centrifugation conditions. Pellet can be decanted, flash-frozen, then stored in -80°C for later use. Cells were then resuspended in 4 mL lysis buffer LB1 (50 mM HEPES, 140 mM NaCl, 1 mM EDTA, 10% glycerol, 0.5% Igepal CA-630, 0.25% Triton X-100, pH to 7.5 using KOH, add 1x protease inhibitor right before use) for 10 min at 4°C, then spun down 2,000 *× g* at 4°C for 3 min. The supernatant was decanted. Cells were then resuspended in 4 mL LB2 (10 mM Tris-HCl pH 8, 200 mM NaCl, 1 mM EDTA, 0.5 mM EGTA, pH to 8.0 using HCl, add 1x protease inhibitor right before use) for 5 min at 4°C, spun down with the same protocol, and the supernatant decanted. Cells were then resuspended in 1.5 mL LB3 (10 mM Tris-HCl pH 8, 100 mM NaCl, 1 mM EDTA, 0.5 mM EGTA, 0.1% Na-Deoxycholate, 0.5% N-lauroylsarcosine, pH to 8.0 using HCl, add 1x protease inhibitor right before use) and transferred to 2 mL Eppendorf tubes for sonication with 50% amplitude, 30 s ON, 30 s OFF for 12 min total time (Fisher 150E Sonic Dismembrator). Sample was spun down with 20,000 *× g* at 4°C for 10 min, and supernatant was transferred to 1.5 mL LB3 in a 15 mL falcon tube. 300 μL of 10% Triton X-100 was added, and the entire solution was well mixed by gentle inversion.

Beads pre-loaded with antibodies were prepared before cell harvesting. 50 μL Protein A beads (Thermo Fisher) were used per IP and transferred to a 2 mL Eppendorf tube on a magnetic stand. Beads were washed twice with blocking buffer BB (0.5% BSA in PBS), then resuspended in 100 μL BB per IP. 4 μL of MRE11 antibody (Novus NB100-142) per IP was added and placed on rotator for 1-2 hours. Right before IP, the 2 mL tube was placed on a magnetic rack and washed 3x with BB, before resuspending in 50 μL BB per EP. 50 μL of beads in BB were transferred to each IP and placed in 4°C rotator for 6+ hours.

Samples were transferred to 2 mL Eppendorf tubes on a magnetic stand, washed 6x with 1 mL RIPA buffer (50 mM HEPES, 500 mM LiCl, 1 mM EDTA, 1% Igepal CA-630, 0.7% Na-Deoxycholate, pH to 7.5 using KOH), then washed 1x with 1 mL TBE buffer (20 mM Tris-HCl pH 7.5, 150 mM NaCl), before decanting. Beads containing ChIP-ed DNA were mixed with 70 μL elution buffer EB (50 mM Tris-HCl pH 8.0, 10 mM EDTA, 1% SDS) and incubated 65°C for 6+ hours. 40 μL of TE buffer was mixed to dilute the SDS, followed by 2 μL of 20 mg/mL RNaseA (New England BioLabs) for 30 min at 37°C. 4 μL of 20 mg/mL Proteinase K (New England BioLabs) was added and incubated for 1 hours at 55°C. The genomic DNA was column purified (Qiagen) and eluted in 35 μL nuclease free water.

Oligo sequences for library preparation are in Table S4. End-repair/A-tailing was performed on 17 μL of ChIPed DNA using NEBNext^®^ Ultra™ II End Repair/dA-Tailing Module (New England BioLabs), followed by ligation (MNase_F/MNase_R) with T4 DNA Ligase (New England BioLabs). 13 cycles of PCR using PE_i5 and PE_i7XX primer pairs were performed for MRE11 ChIP samples to amplify sequencing libraries. Samples were pooled, quantified with QuBit (Thermo), Bioanalyzer (Agilent) and qPCR (BioRad).

Cell line samples were sequenced on a NextSeq 500 (Illumina) using paired 2×36bp reads. Mouse liver samples were sequenced on a DNBSEQ PE100 (BGI) using paired 2×50bp reads. All ChIP-seq raw reads in FASTQ format and processed alignments in BAM format are uploaded to Sequence Read Archive (SRA) under BioProject accession PRJNA801688.

Reads were demultiplexed after sequencing using bcl2fastq. Paired-end reads were aligned to hg38, hg19, or mm10 using bowtie2. To ensure fair comparison between DISCOVER-Seq+ (with DNA-PKcs inhibitor) and DISCOVER-Seq (without inhibitor), equal numbers of sequencing reads were obtained by subsetting for each set of samples. Samtools was used to filtered for mapping quality >= 25, remove singleton reads, convert to BAM format, remove potential PCR duplicates, and index BAM-formatted output files. The software that coordinates these steps as well as performs subsequent analyses are open source (https://github.com/rogerzou/DSeqPlus).

BLENDER (Wienert et al., 2019) (https://github.com/staciawyman/blender) was used to determine Cas9 off-target sites, outputting a curated list of all off-target sites with corresponding visualization. A more sensitive cutoff threshold of 2 (-c 2) was used for all samples except a threshold of 3 (-c 3) for the merged *PCSK9* samples.

### CRISPR-Cas9 or Cas12a editing of primary human T cells

Engineered T cells expressing a TP53 R175H:HLA-A*02:01-specific TCR under control of a EF1-alpha promoter were generated via CRISPR-Cas-mediated homology directed repair (HDR) electroporation as follows. Nucleotide sequences of the TCR of interest, promoter, and homology arms for the *TRAC* gene locus were generated by de-novo gene synthesis (GeneArt). A 4699 bp HDR template dsDNA (HDRT) was generated by amplification from a plasmid template using the Q5 High-Fidelity 2X Master Mix (New England BioLabs) with primers containing truncated Cas9 target sequences (IDT) [PMID: 31819258]. Amplicon DNA was purified with AMPure beads (Beckman Coulter), eluted in water, and quantified. Purified PCR products were analyzed by agarose gel electrophoresis to assess correct amplicon size and purity. T cells were isolated by negative selection using immunomagnetic cell separation (EasySep Human T Cell Isolation Kit) from cryopreserved healthy donor peripheral blood mononuclear cells collected via leukapheresis. Purified CD3+ T cells were activated with Dynabeads Human T-Activator CD3/CD28 (ThermoFisher) at a 1:2 bead-to-cell ratio in RPMI-1640 (ATCC) supplemented with 10% fetal bovine serum (HyClone Defined), 100 units/mL Penicillin (Gibco), 100 ug/mL Streptomycin (Gibco), 100 IU/mL recombinant human IL-2 (Proleukin, Prometheus Laboratories) and 5 ng/mL recombinant human IL-7 (BioLegend) at 37C, 5% CO_2_. After 48 hours and prior to electroporation, beads were removed with a magnet. Cas9 ribonucleoprotein (RNP) targeting *TRAC* (AGAGTCTCTCAGCTGGTACA) or Cpf1 (Cas12a) RNP targeting a juxtaposed nucleotide sequence in *TRAC* (GAGTCTCTCAGCTGGTACAC) were assembled by mixing the appropriate sgRNA (IDT) with either Alt-R S.p. Cas9 nuclease V3 (IDT) or Alt-R A.s. Cas12a (Cpf1) Ultra nuclease (IDT) and matching ssDNA Electroporation Enhancer (IDT) and incubating the mixture at room temperature for 15 minutes. RNPs were mixed with 0.5 μg of the same HDRT and incubated for 5 minutes at room temperature. To edit activated T cells, 20 μL of T cells were resuspended in P3 buffer at 5 × 10^7^ cells/mL (Lonza) and added to the electroporation mixture. Electroporation was performed with a 4D-Nucleofector X Unit (Lonza) in 16-well cuvettes using pulse code EH115. After electroporation, T cells were recovered by immediately adding 80 uL of warm, cytokine-free T cell media to the cuvettes and incubation at 37 °C for 15 minutes. Then, T cells were diluted in T cell growth media containing 100 IU/mL recombinant human IL-2 and 5 ng/mL recombinant human IL-7 in the presence of 1 μM Ku-60648 or vehicle (DMSO) and incubated for 12 hours at 37C, 5% CO_2_. A fraction of electroporated T cells for each condition was maintained in T cell growth media with human IL-2 and IL-7 for 7 days prior to analysis for gene editing rates and surface expression of TP53 R175H:HLA-A*02:01-specific TCR by flow cytometry.

### High throughput sequencing of genomic DNA samples

Genomic DNA was extracted using the DNeasy Blood & Tissue Kit (Qiagen 69504) following manufacturer instructions. Approximately 1 million cells were used from cell lines and iPSCs. Approximately 10-20 μL of mouse liver cell suspension was used out of 1.5 mL total, and the genome extraction protocol included the Buffer ATL step for tissue lysis.

Genomic DNA samples were amplified with PCR using Q5 Hot Start High-Fidelity 2X Master Mix (New England BioLabs M0494). Primer pairs for all sequences are listed in Table S3. For example, the primer set for amplifying around the *FANCF site 2* on-target site is NGS_Fs2_ON_F and NGS_ Fs2_ON_R. After amplicon PCR, cleanup was performed using 1.4x AMPure XP (Beckman Coulter A63881) following the manufacturer’s instructions. Dual-indexing PCR was performed using KAPA HiFi HotStart ReadyMix (Roche 07958935001) and PCR cleanup was performed using 1x AMPure XP. Samples were quantified using QuBit (Thermo Fisher Scientific), pooled, diluted, and loaded onto a MiSeq (Illumina). Sequencing was performed with the following number of cycles “151 | 8 | 8 | 151” with the paired-end Nextera sequencing protocol.

Sequencing reads were either demultiplexed automatically using MiSeq Reporter (Illumina) or with a custom Python script to individual FASTQ files. For indel calling, sequencing reads were scanned for exact matches to two 20-bp sequences that flank +/-20 bp from the ends of the target sequence. If no exact matches were found, the read was excluded from analysis. After additional filtering for an average quality score > 20, an indel is defined as a sequence that differs in length from the reference length.

