## Supplementary Materials for "Improving the sensitivity of in vivo CRISPR off-target detection with DISCOVER-Seq+"

1 Department of Biomedical Engineering, Johns Hopkins University School of Medicine, Baltimore, Maryland, USA
2 Department of Biophysics and Biophysical Chemistry, Johns Hopkins University School of Medicine, Baltimore, Maryland, USA
3 Department of Biophysics, Johns Hopkins University, Baltimore, Maryland, USA
4 Department of Medicine, Johns Hopkins University School of Medicine, Baltimore, Maryland, USA
5 Division of Rheumatology, Department of Medicine, Johns Hopkins University School of Medicine, Baltimore, MD, USA.
6 Ludwig Center, Sidney Kimmel Comprehensive Cancer Center, Johns Hopkins University School of Medicine, Baltimore, MD, USA.
7 Lustgarten Pancreatic Cancer Research Laboratory, Sidney Kimmel Comprehensive Cancer Center, Johns Hopkins University School of Medicine, Baltimore, MD, USA.
8 Howard Hughes Medical Institute, Chevy Chase, MD, USA.
9 Equal Contribution

Supplementary Figures 1-2
Supplementary Table 1

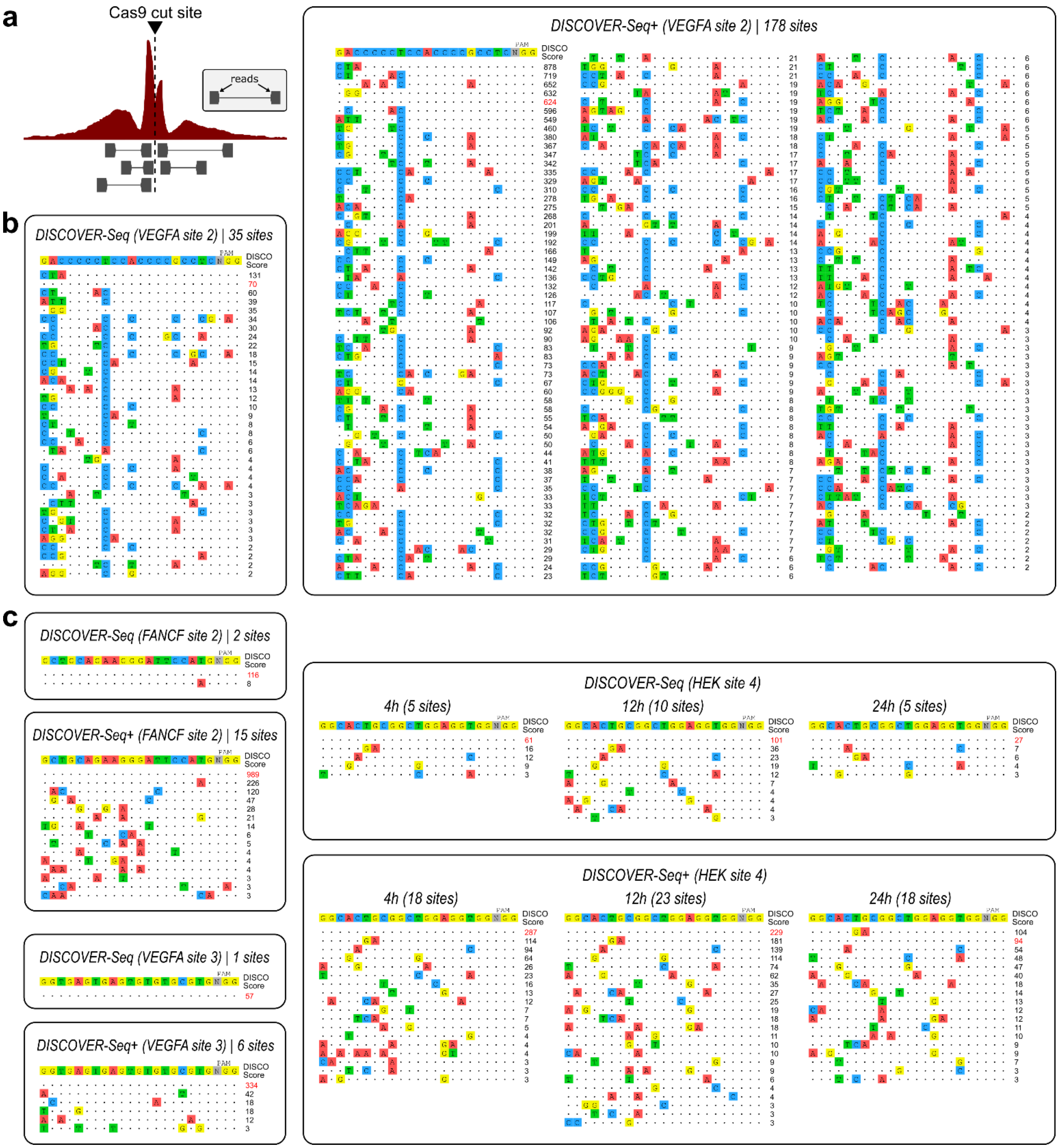


**Figure S1: Lists of off-target sites discovered by MRE11 ChIP-seq and improved with DNA-PKcs inhibition**

**a,** Illustration of MRE11 ChIP-seq enrichment at a specific target site, visualized as a histogram of base pair coverage from sequencing reads along the genome. Only the two ends of each DNA fragment are sequenced.

**b,** *VEGFA site 2* Cas9 target sites detected using DISCOVER-Seq (left) versus DISCOVER-Seq+ (right) in K562 cells.

**c,** *FANCF site 2, VEGFA site 3, or HEK site 4* Cas9 target sites detected using DISCOVER-Seq versus DISCOVER-Seq+ in K562 or HEK293T cells, at 4 h, 12 h, or 24 h after Cas9 delivery.

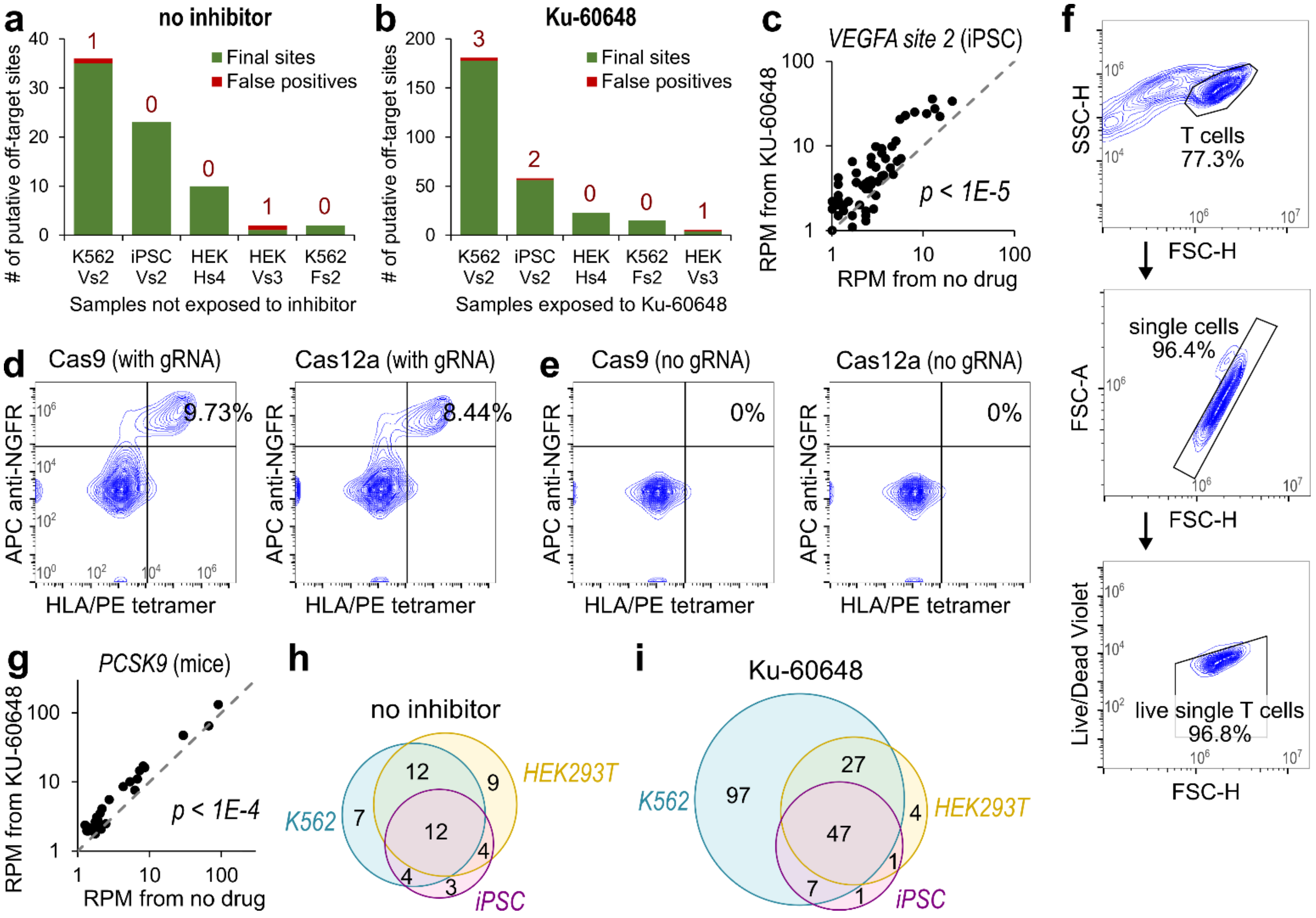


**Figure S2: Validation of DISCOVER-Seq+ in diverse applications**

**a-b,** Plot of the total number of initial off-target sites, for samples (a) without inhibitor and (b) with Ku-60648. Sites labeled as false positive, because they were also reported in corresponding negative control samples without Cas9, are colored red with the number of sites indicated above each bar. For x-axis, first row is cell type, second row is gRNA target (Vs2: VEGFA site 2, Hs4: HEK site 4, Vs3: VEGFA site 3, Fs2: FANCF site 2).

**c,** Plot of MRE11 ChIP-seq enrichment at all DISCOVER-Seq+ detected target sites within a 1.5 kb window for Cas9 targeting *VEGFA site 2* in WTC-11 iPSCs. Each point in the plot (55 total) corresponds to a putative target site. MRE11 enrichment from DISCOVER-Seq+ data (y-axis) versus DISCOVER-Seq data (x-axis) was significantly different ($p < 1E-5$), determined using the two-sided Wilcoxon rank sum test.

**d-e,** (d) Flow cytometry contour plots of T cell populations at day 7 following CRISPR editing with Cas9 (left) or Cas12a (right). X-axis gates by T cell binding to a phycoerythrin (PE)-conjugated HLA-A*02:p53 R175H peptide tetramer complex specific for the transgenic TCR (tgTCR). Y-axis gates by allophycocyanin (APC)-conjugated antibody against NGFR (CD271), introduced as part of the CRISPR HDRT as an editing control. Cells positive for both markers represent successful cancer-specific tgTCR integration. 9.7% of T cells are positive for both markers using Cas9 compared to 8.4% with Cas12a. (e) Same, but for negative control samples without gRNA.

**f,** Gating strategy for flow cytometric analysis of live single T cells, used for panels d-e.

**g,** Same as panel c, for Cas9 targeting *PCSK9* in the liver of mice, with 30 total target sites. MRE11 enrichment from DISCOVER-Seq+ was significantly different ($p < 1E-4$), using the two-sided Wilcoxon rank sum test.

**h-i,** Venn diagram illustrating overlap in the *VEGFA site 2* sites identified by (h) DISCOVER-Seq or (i) DISCOVER-Seq+ in K562 cells (blue), HEK293T cells (yellow), and WTC-11 iPSCs (purple).

| | | Discovery of CRISPR off-targets has successfully been shown in | | | | | | |
|---|---|---|---|---|---|---|---|---|
| off-target method | reference | DNA only | transformed cells | primary cells | animal tissues | sensitivity | false positives | reporter required |
| DISCOVER-Seq+ | this manuscript | - | + | + | + | ++ | - | - |
| DISCOVER-Seq | Wienert et al., 2019 | - | + | + | + | + | - | - |
| BLESS/BLISS | Yan et al., 2017 | - | + | - | - | ++ | + | - |
| GUIDE-tag | Liang et al., 2022 | - | - | - | + | ++ | - | + |
| GUIDE-seq | Tsai et al., 2015 | - | + | - | - | ++ | - | + |
| CIRCLE-seq | Tsai et al., 2017 | + | - | - | - | +++ | +++ | - |
| SITE-seq | Cameron et al., 2017 | + | - | - | - | +++ | +++ | - |
| Digenome-seq | Kim et al., 2015 | + | - | - | - | +++ | +++ | - |

**Supplementary Table 1:** Comparison of DISCOVER-Seq+ to other methods for detecting off-target sites. Part of the table is derived from Wienert et al., 2019.
